# Evaluation of Deep Neural Network Models for Instance Segmentation of Lumbar Spine MRI

**DOI:** 10.1101/2024.04.02.587810

**Authors:** Jiasong Chen, Linchen Qian, Linhai Ma, Timur Urakov, Weiyong Gu, Liang Liang

**Affiliations:** Department of Computer Science, University of Miami, Coral Gables, FL; Department of Neurological Surgery, University of Miami, Coral Gables, FL; Department of Mechanical and Aerospace Engineering, University of Miami, Coral Gables, FL

**Author notes:** For correspondence: Liang Liang, Ph.D., Department of Computer Science, University of Miami, Ungar Building, Room 330K, Coral Gables, FL, 33146.

**Keywords:** Lumbar spine MRI, Medical image instance segmentation, Data augmentation

## Abstract

Intervertebral disc disease, a prevalent ailment, frequently leads to intermittent or persistent low back pain, and diagnosing and assessing of this disease rely on accurate measurement of vertebral bone and intervertebral disc geometries from lumbar MR images. Deep neural network (DNN) models may assist clinicians with more efficient image segmentation of individual instances (discs and vertebrae) of the lumbar spine in an automated way, which is termed as instance image segmentation. In this work, we evaluated 15 existing DNN models for lumbar spine MR image segmentation. We introduced a new data augmentation technique to create synthetic yet realistic MR image dataset, named SSMSpine, which is made publicly available. The 15 image segmentation models are evaluated on our private in-house dataset and the public SSMSpine dataset, using two metrics, Dice Similarity Coefficient and 95% Hausdorff Distance. The SSMSpine dataset are available at https://github.com/jiasongchen/SSMSpine.

## 1. Introduction

The intervertebral discs in humans can undergo a profound degenerative process as early in the adolescence (Cox et al., 2014; Kos et al., 2019), which can be accompanied by facet arthropathy and hypertrophy. This degeneration can manifest as various conditions, including discogenic low back pain, disc herniation, spinal stenosis, and spondylolisthesis, which may necessitate the implementation of surgical or non-surgical interventions aimed at alleviating pain and restoring normal functionality. Magnetic resonance imaging (MRI) is the most widely used technique for specifically quantifying intervertebral discs degeneration (IDD) by assessing changes in disc geometry deformation and signal strength degradation (Mallio et al., 2022; Roberts et al., 2021; Tamagawa et al., 2022). The information derived from imaging data is of utmost importance for medical professionals in terms of both diagnosing medical conditions and planning appropriate treatments. Furthermore, this information serves as a critical foundation for developing patient-specific computational models, which hold the potential to mature over time and eventually enable accurate predictions of treatment outcomes within clinical settings. Presently, the process of geometry reconstruction, signal measurements, and grading from magnetic resonance (MR) images heavily relies on manual annotation. However, this process is not only time-consuming but also vulnerable to human bias. Consequently, there is an urgent need for automated MR image analysis methods to address these challenges.

In medical imaging, semantic/instance image segmentation, which divides the images into distinct sections at the pixel level so that each pixel belongs to a specific region, has the potential to be carried out through automated techniques (Galbusera et al., 2019). The traditional methods, such as watershed and level set, have demonstrated satisfactory performance in medical image segmentation tasks. The watershed method treats an image as a topological map where intensity represents the altitude of the pixels. The watershed segmentation is determined by the watershed lines on a topographic surface (Chevrefils et al., 2007; Huang and Chen, 2004). The level set method performs image segmentation by utilizing dynamic variational boundaries (Huang et al., 2013). However, the traditional method suffers from the clinical variation of different patients and the noise effect of different medical imaging equipment, problems like the over-segmentation and time-consuming consist (Li et al., 2007).

Since the increasingly vast amount of medical imaging data and computational resources have become available, machine learning (ML) methods, especially deep neural network techniques, show superior performance than traditional methods. Convolutional neural network (CNN) has a significant edge over its predecessors in that it possesses the capability to recognize essential components/features without requiring any human intervention (Suganyadevi et al., 2022). CNNs are specifically designed to effectively utilize spatial and configural information by accepting 2D or 3D images as input. This approach helps to prevent the loss or disruption of structural and configural information in medical images (Shen et al., 2017).Various deep CNNs, including UNet++ (Zhou et al., 2018), Attention U-Net (Oktay et al., 2018), MultiResUNet (Ibtehaz and Rahman, 2020) and UNeXt (Valanarasu and Patel, 2022) have been proposed for image segmentation for different medical imaging modalities and different organs (e.g. heart (Cao et al., 2023; Gao et al., 2021; Huang et al., 2023), lung (Zhou et al., 2018), brain (Hatamizadeh et al., 2022, 2021; Hu et al., 2022; Valanarasu et al., 2021), pancreas (Oktay et al., 2018), gland (Valanarasu et al., 2021; Wang et al., 2022), spine (Sekuboyina et al., 2018; Wang et al., 2023), retina blood vessels (Moccia et al., 2018; Soomro et al., 2019), aorta (Berhane et al., 2020; Noothout et al., 2018; Pepe et al., 2020), etc). Although these methods have achieved promising performance, there are still some limitations in a more complex context coping with long-range dependency explicitly due to the intrinsic locality of convolutions.

Recently, Transformer, an ML technique, has shown exceptional performance not only on natural language processing (NLP) challenges like machine translation (Vaswani et al., 2017), but also image analysis tasks including image classification (Shamshad et al., 2023) and segmentation (Chen et al., 2021; Hatamizadeh et al., 2022, 2021; Liu et al., 2021; Wang et al., 2022). Various variations of Transformer models have demonstrated that the global information perceived by the self-attention operations is beneficial in medical imaging tasks. TransUNet was the first Transformer-based network specifically for medical image segmentation on the synapse multi-organ segmentation dataset (Chen et al., 2021). Wang et al. (2022) substituted the original skip connection scheme of U-Net with the proposed UCTransNet that includes a multi-scale Channel Cross fusion Transformer and a Channel-wise Cross-Attention and tested the network on the gland segmentation dataset (Sirinukunwattana et al., 2017) and synapse multi-organ segmentation dataset (Landman et al., 2015). Hatamizadeh et al., (2021, 2022) proposed both UNETR and Swin UNETR for 3D medical imaging segmentation. UNETR utilizes a U-shape network with a vision Transformer as the encoder and a CNN-based decoder. Swin UNETR is constructed by replacing the vision transformer encoder in UNETR architecture with the Swin Transformer encoder. Feng et al. (2022) proposed SLT-Net to utilize CSwin Transformer (Dong et al., 2022) as the encoder for feature extraction and the multi-scale context Transformer as the skip connection for skin lesion segmentation. Swin-Unet adopted Swin Transformer (Liu et al., 2021) with shifted windows as encoder and a symmetric Swin Transformer-based decoder with patch expanding layer as decoder for multi-organ segmentation task (Cao et al., 2023). Pu et al., (2023) proposed a semi-supervised learning framework with Inception-SwinUnet adopting convolution and sliding window attention in different channels for vessel segmentation on small amount of labeled data. Besides the self-attention mechanism, position embeddings are another crucial component of Transformer models. Regarding changing the order of the input, a Transformer model is invariant (Vaswani et al., 2017) without position embeddings. However, since text data inherently has a sequential structure, the absence of position information results in the ambiguous or undefined meaning of a sentence (Dufter et al., 2022). For image segmentation, usually, an image patch is treated as a token, and Transformers process the entire input sequence of tokens in parallel. With position embeddings, a Transformer would be able to differentiate between image patches with similar content that appear in different positions in the input image, which is beneficial for image segmentation applications. A variety of different methods may be used to incorporate the position information into Transformer models. Absolute position encoding and relative position encoding are two main categories to encode a token’s position information. Vaswani firstly introduced absolute and relative position embedding in the vanilla Transformer model (Vaswani et al., 2017). Shaw extended the self-attention mechanism with the capacity of effectively incorporating the representation of relative position (Shaw et al., 2018). Valanarasu et al. (2021) proposed a gated position-sensitive axial attention mechanism to cope the difficulty in learning position encoding for the images.

Specifically for lumbar spine research, instance segmentation of MR images is preferred, which not only determines whether or not a pixel belongs to a disc, but also labeling the precise instance to which it belongs (Galbusera et al., 2019). In recent years, most instance segmentation methods for spine image segmentation are based on CNNs-only networks, and only a few Transformer-based networks are employed. For example, Kuang et al. (2020) built an unsupervised segmentation network for spine image segmentation using the rule-based region of interest (ROI) detection, a voting mechanism accompanied by a CNN network. Sekuboyina et al. (2018) proposed a dual branch fully convolutional network that take advantages of both low-resolution attention information on two-dimensional sagittal slices and high-resolution segmentation context on three-dimensional patches for effective segmentation of the vertebrae. MLKCA-Unet incorporates multi-scale large-kernel convolution and convolutional block attention into the U-net architecture for efficient feature extraction in spine MRI segmentation (Wang et al., 2023). Pang et al. (2022) introduced a mixed-supervised segmentation network and it was trained on a strongly supervised dataset with full segmentation labels and a weakly-supervised dataset with only key points. BianqueNet combined new modules with a modified deeplabv3+ network (Chen et al., 2018), which includes a Swin Transformer-skip connection module, for segmentation of lumbar intervertebral disc degeneration related regions (Zheng et al., 2022).

It was shown that the Transformer-based models only perform effectively when trained on large-scale datasets since the lack of inductive bias (Dosovitskiy et al., 2021). The utilization of Transformer-based networks for medical imaging tasks poses a challenge due to the limited availability of labeled images in medical datasets. Obtaining well-annotated medical imaging datasets presents significantly greater challenges compared to curating traditional computer vision datasets. Dealing with expensive imaging equipment, complex image acquisition pipelines, expert annotation requirements, and privacy concerns are all part of the problematic issues (Litjens et al., 2017). This scarcity hampers the effective application of Transformer-based models in the medical domain. In such scenarios, the adoption of suitable and feasible data augmentation techniques becomes crucial prior to model training. These techniques can help to increase the effective size of the medical image dataset and improve the performance of the Transformer-based model.

In this study, we evaluated 15 DNN modes for lumbar spinal MRI instance segmentation. For this purpose, we developed a novel data synthesis method based on statistical shape model (SSM) and biomechanics. This SSM-biomechanics-based data synthesis method generates lumbar spine images with large and plausible deformations, which can be used for model training and evaluation.

## 2. Review of Transformer-based DNN models for Lumbar Image Segmentation

### 2.1. Transformer-based Networks for the segmentation of non-spine medical images

Transformer-based networks have shown promising performance on medical image segmentation because of their ability to capture long-range dependencies. Comparably, the inductive bias of CNN networks benefits from its local connectivity and parameter sharing property. Therefore, many networks, combining CNN and Transformers and leveraging both benefits, have been proposed in the past few years. TransUNet (Chen et al., 2021) firstly combines CNNs and Transformer in a cascaded manner in its encoder for medical imaging tasks, in which the low-level features are collected from the CNNs and then fed to the Transformer to capture global interactions. Other designs of CNN and Transformer networks, including UNETR (Hatamizadeh et al., 2021), UTNet (Gao et al., 2021), UCTransNet (Wang et al., 2022), Swin UNETR (Hatamizadeh et al., 2022) and MedT (Valanarasu et al., 2021) have shown better segmentation performances in different medical image modalities, compared to CNN-only networks. UNETR replaces the encoder of a UNet with Transformer layers with 1D learnable positional embedding, and its decoder only has convolution layers (Hatamizadeh et al., 2021). UTNet inserts Transformer layers into a Unet in a sequential manner: a convolution layer followed by a Transformer layer with learnable relative position embedding (Gao et al., 2021). UCTransNet embeds Transformer layers into the skip-connections of a Unet with fully learnable absolute position embedding (Wang et al., 2022). Swin UNETR replaces the Transformer in UNETR with Swin-Transformer (Hatamizadeh et al., 2022). Medical Transformer (MedT) uses gated-axial Transformer layers in the encoder of a Unet (Valanarasu et al., 2021). HSNet used PVTv2 (Wang et al., 2022) as encoder and a dual-branch structure which Transformer branch and CNN branch fused by element-wise product as decoder for polyp segmentation (Zhang et al., 2022).

### 2.2. Transformer-based Networks for the segmentation of spine images

To the best of our knowledge, there are only a few Transformer-based networks (You et al., 2022; Tao et al., 2022) specifically designed for lumbar spine image segmentation, including EG-Trans3DUNet (You et al., 2022), Spine-transformers (Tao et al., 2022), APSegmenter (Zhang et al., 2022), and BianqueNet (Zheng et al., 2022). However, it is worth noting that most of these networks were developed for the segmentation of vertebral bodies using CT modality, which may not be directly applicable to the segmentation of the intervertebral discs in order to study disc degeneration. Amony those networks, only the code of BianqueNet is publicly available.

EG-Trans3DUNet combines two vision transformer branches to handle both local patches and resized global spinal CT images (You et al., 2022), and it merges edge characteristics and semantic features generated by a CNN-based edge detection block. Spine-transformers was designed for spinal CT image segmentation with a two-stage pipeline to handle arbitrary Field-Of-View input images (Tao et al., 2022). In its first stage, a Transformer with a CNN backbone is utilized as a 3D object detector to locate individual vertebrae, and then the input image is cropped into regions of individual vertebrae. In its second stage, a multi-task encoder-decoder CNN network is applied to each cropped region to segment the vertebra. The source code of the second stage is not publicly available. APSegmenter (Zhang et al., 2022) combines a ViT-style Transformer with a mask Transformer to segment spine X-ray images, and an adaptive postprocessing is applied to further refine the result. BianqueNet (Zheng et al., 2022) employed a resnet101 network to perform feature extraction, followed by upsampling using the Swin Transformer-skip connection module and a double upsampling operations. It also used a multi-scale feature fusion module to generate the segmentation of regions associated with intervertebral disc degeneration.

In addition to the segmentation of vertebral bodies, the segmentation of intervertebral discs (IVDs) is vital for lumbar spinal disease diagnosis and treatment. Since the water content in IVDs cannot be revealed on CT images, currently, MRI is the gold standard imaging modality for the evaluation of IVD pathologies (Kirnaz et al., 2022). Our study aims to explore the benefit of combining CNN and transformer for instance segmentation of lumbar spine MR images.

### 2.3. Self-Attention in Transformer

As opposed to convolutional operations, the self-attention mechanism within a Transformer network has the fundamental advantage of effectively capturing global features and long-range contextual dependency. It uses the Key, Query and Value vectors to better describe the features’ connections. Nonetheless, because of the inherited properties of self-attention, it doesn’t retrieve the position information on its own, which is important for instance segmentation. One of the best ways to tackle this problem is to use a well-designed position embedding mechanism to inject the position relationships into the self-attention calculation.

#### 2.3.1. The classic self-attention mechanism with additive position embedding

The plain Transformer is constructed with the multi-head self-attention modules (MHSA), which enable Transformer to capture and utilize more accurate and detailed spatial information (Vaswani et al., 2017). Given an input token set (e.g., image patches) *X*, three individual linear transformations (*W*_*Q*_, *W*_*K*_, *W*_*V*_) are applied to *X* to generate query embedding (*Q*), key embedding (*K*), and value embedding (*V*). Then, the self-attention score, *Attn*, is calculated as a scale-product of these three embedding as following:

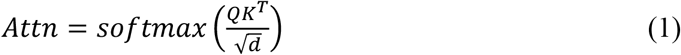

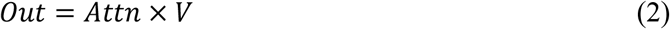

In the above equations, *Q* = (*X* + *P*)*W*_*Q*_, *K* = (*X* + *P*)*W*_*K*_, *V* = (*X* + *P*)*W*_*V*_. *P* is the encoded position, and *d* is the dimension of embedding in each head. *Out* is the final output of the self-attention module.

#### 2.3.2. Position Embedding

Without using any position embedding, the self-attention mechanism in Eq.(1) is permutation-invariant and cannot distinguish tokens (e.g., image patches) at different spatial locations. Therefore, it is essential to design efficient position embedding cooperating with the self-attention mechanism.

The attention matrix in the classic self-attention Eq.(1) can be decomposed into three terms:

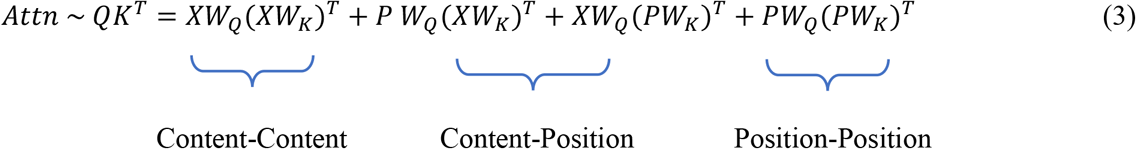

Therefore, the attention considers three interactions/correlations among tokens: *XW*_*Q*_ (*XW*_*K*_)^*T*^ for content to content interaction, *P W*_*Q*_ (*XW*_*K*_)^*T*^ + *XW*_*Q*_ (*PW*_*K*_)^*T*^ for interaction between content and position, and *PW*_*Q*_ (*PW*_*K*_)^*T*^ for position to position interaction. As the instance segmentation task is location-specific, a well-designed interaction between content and position could improve self-attention ability to utilize both content and position information to accomplish the instance segmentation task.

Generally, there are mainly two steps to define the position embedding. The first step is defining the position function or distance function, which is used for encoding the position information of input tokens. There are plenty of position functions such as index function, Euclidean distance, and sinusoidal functions etc., The second step is defining methods to incorporate the encoded position information into self-attention.

Absolute position embedding and relative position embedding are two main position representation methods to incorporate the position information into input tokens. Absolute position embedding encodes the absolute positions of each input tokens as individual encoding vectors, and relative position embedding focus on the relative positional relationships of pairwise input tokens (Lin et al., 2022; Wu et al., 2021). In the vanilla Transformer designed for NLP (Vaswani et al., 2017), it used a combination of absolute and relative position embedding to add position information to the tokens. It is inconclusive that if relative position embedding is better or worse than absolute position embedding, and the answer seems to be dependent on specific applications (Dufter et al., 2022; Huang et al., 2020; Shaw et al., 2018; Wu et al., 2021). Relative position encoding benefits from capturing the details of relative distance/direction and is invariant to tokens’ shifting. The intuition is that, in the self-attention mechanism, the pairwise positional relationship (both in terms of direction and distance) between input elements might be more advantageous than absolute position of individual elements (Lin et al., 2022). In such a case, position information in Transformer is an extensive research area, and various relative position encodings have been proposed for medical imaging segmentation (Dosovitskiy et al., 2021). For example, UTNet proposed the 2-dimensional relative position encoding by adding relative height and width information (Gao et al., 2021). MedT updated self-attention mechanism with position encoding along the width axis with the inspiration of axial attention (Valanarasu et al., 2021; Wang et al., 2020; Zhang and Zhang, 2022). The Parameter-Efficient Transformer added a trainable position vector to the input to encode relative distances (Hu et al., 2022). In this work, we propose a novel relative position embedding method for segmentation performance improvement.

#### 2.3.3. Image Self-Attention in the Existing Image Segmentation Models

We compared instance segmentation performances of 15 representative image segmentation models, including 11 Transformer-based models and 4 CNN-only models.

Table 1 summarizes the image self-attention mechanisms in the existing 11 Transformer-based models. TransUnet and UNETR add the position embedding directly into the input patches (Chen et al., 2021; Hatamizadeh et al., 2021). Swin-Unet, Swin UNETR, and Inception-SwinUnet incorporate the position embedding into attention score instead of input tokens. (Cao et al., 2023; Hatamizadeh et al., 2022; Pu et al., 2023). BianqueNet utilized the position embedding within Swin-Transformer (Zheng et al., 2022). According to the source code, SLT-Net introduces a lepe distance to represent the position bias (embedding), which is directly added into the output matrix (Feng et al., 2022). MedT proposed a gated position-sensitive axial attention mechanism where four learnable gates (*G*) control the amount of position embedding contained in key (*K*), query (*Q*) and value (*V*) embeddings (Valanarasu et al., 2021). UTNet introduced the 2-dimensional relative position encoding by adding relative position logits along height and width dimensions(*R*_*height*_, *R*_*width*_) into the key embedding (Gao et al., 2021). HSNet and UCTransNet do not include position embedding in their models (Zhang et al., 2022; Wang et al., 2022)

**Table 1.** Image Self-Attention in the attention/Transformer-based segmentation models.

| Model | Self-Attention |
| --- | --- |
| Swin UNETR | $\text{softmax}\left(\frac{XW_Q(XW_K)^T}{\sqrt{d}} + R\right)(XW_V)$<br>Note: $R$ is called relative position bias in the reference |
| SLT-Net | $\text{softmax}\left(\frac{XW_Q(XW_K)^T}{\sqrt{d}}\right)(XW_V) + \text{lepe}(XW_V)$<br>Note: $\text{lepe}$ is a convolution kernel |
| UNETR | $\text{softmax}\left(\frac{(X+P)W_Q((X+P)W_K)^T}{\sqrt{d}}\right)(X+P)W_V$ |
| Inception-SwinUnet | $\text{softmax}\left(\frac{XW_Q(XW_K)^T}{\sqrt{d}} + R\right)(XW_V)$<br>Note: $R$ is called relative position bias in the reference |
| HSNet | $\text{softmax}\left(\frac{XW_Q(XW_K)^T}{\sqrt{d}}\right)(XW_V)$ |
| Swin-Unet | $\text{softmax}\left(\frac{XW_Q(XW_K)^T}{\sqrt{d}} + R\right)(XW_V)$<br>Note: $R$ is called relative position bias in the reference |
| TransUNet | $\text{softmax}\left(\frac{(X+P)W_Q((X+P)W_K)^T}{\sqrt{d}}\right)(X+P)W_V$ |
| MedT | $\text{softmax}\left(XW_Q(XW_K)^T + G_QXW_Q(R_Q)^T + G_KXW_K(R_K)^T\right)(G_{V_1}XW_V + G_{V_2}R_V)$<br>Note: $G_Q, G_K, G_{V_1}, G_{V_2}$ are learnable parameters for gating mechanism.<br>$R_Q, R_K, R_V$ are the position bias for query, key and value |
| UTNet | $\text{softmax}\left(\frac{XW_Q(XW_K + R_{width} + R_{height})^T}{\sqrt{d}}\right)XW_V$<br>Note: $R_{width}$ and $R_{height}$ are the relative height and width information |
| UCTransNet | $\text{softmax}\left(\frac{X_cW_Q(X_cW_K)^T}{\sqrt{d}}\right)(X_cW_V)$<br>Note: $X_c$ is composed of image channels instead of image patches. |
| BianqueNet | $\text{softmax}\left(\frac{XW_Q(XW_K)^T}{\sqrt{d}} + R\right)(XW_V)$<br>Note: $R$ is called relative position bias in the reference |

## 3. Methods

### 3.1. Novel data augmentation/synthesis method based on SSM and biomechanics

For medical image data augmentation, elastic deformation is often used for nonlinear deformation of the images to increase diversity of training data (Ronneberger et al., 2015). Briefly, the input space is discretized by a grid, and a random displacement field on the grid is generated by sampling from a normal distribution with standard deviation equal to *σ* × grid resolution (i.e., the size of a grid cell). The parameter *σ* determines deformation magnitude. To ensure a large deformation with diffeomorphism, the grid needs to be coarser than the input size (i.e., 512 ×512). In this study, we applied two successive elastic deformations to each training image, with grid sizes of 9 × 9 and 17 × 17. As shown in Figure 1, when the deformation parameter *σ* is larger than 0.5, the generated images and spine shapes are highly unrealistic.

**Fig. 1.**
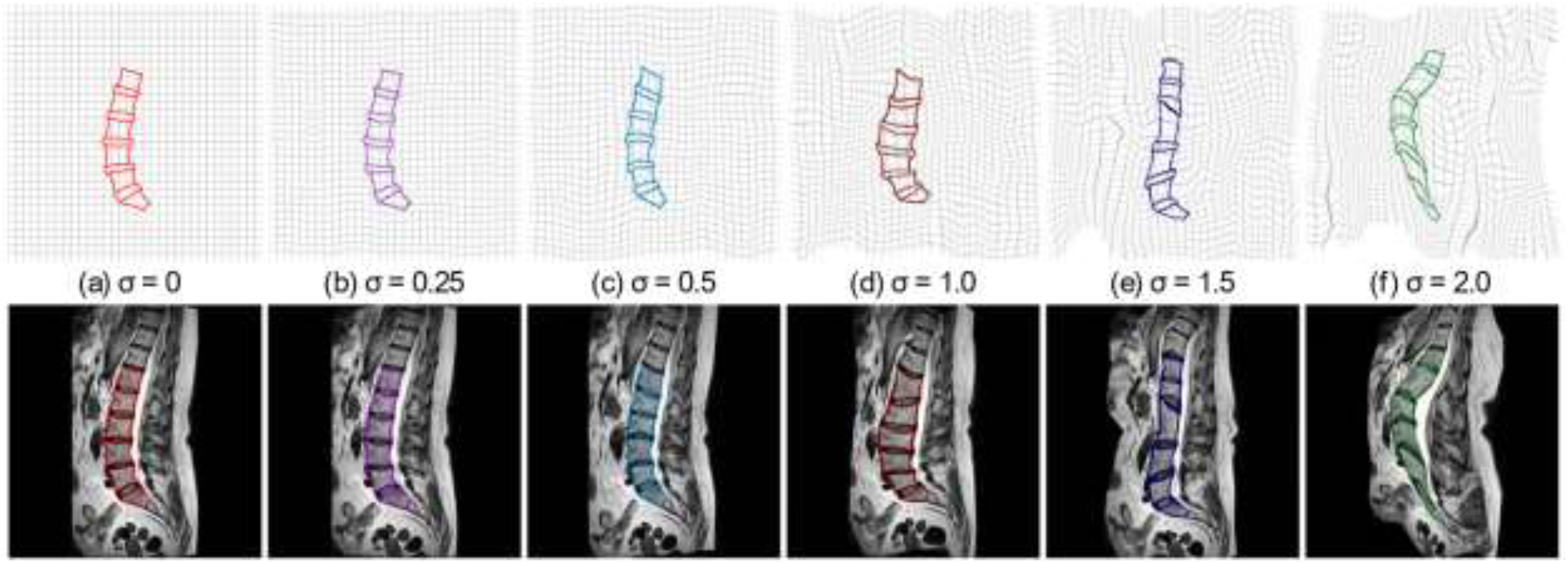
Data augmentation/synthesis examples using elastic deformation with sigma from *σ* to 2.0.

In this work, we developed a new method to synthesize lumbar spine MR images suitable for model training and evaluation. First, we built a statistical shape model (SSM) of lumbar spine shapes (i.e., contours of discs and vertebrae) in a dataset set, and the SSM represents the probability distribution of lumbar spine shapes. We refer the reader to the reference papers (Ambellan et al., 2019; Sarkalkan et al., 2014; Hufnagel et al., 2007; Davies et al., 2003; Cootes et al., 1995) for the details of constructing an SSM. By sampling from the SSM, different lumbar spine shapes can be generated, and each generated lumbar spine shape could be considered from a virtual patient. We note that the SSM technique has been used to generate virtual but realistic patient geometries in many applications, such as generating aortic aneurysm geometries (Liang et al., 2017; van Veldhuizen et al., 2022; Wiputra et al., 2023). Given a lumbar spine shape, if a lumbar spine MR image can be generated and consistent with the shape, then we will have a new sample with ground-truth. For this purpose, we developed a biomechanics-based method to generate a lumbar spine MR image *Ĩ* from a lumbar spine shape 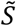 by using a reference image *I* with its ground-truth shape *S*. Intuitively speaking, a nonlinear spatial transform from the shape *S* to the shape 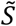 is determined by using biomechanics principles, and then *Ĩ* is obtained by applying the spatial transform to *I*. The generated images are visually plausible, as shown in Figure 2.

**Fig. 2.**
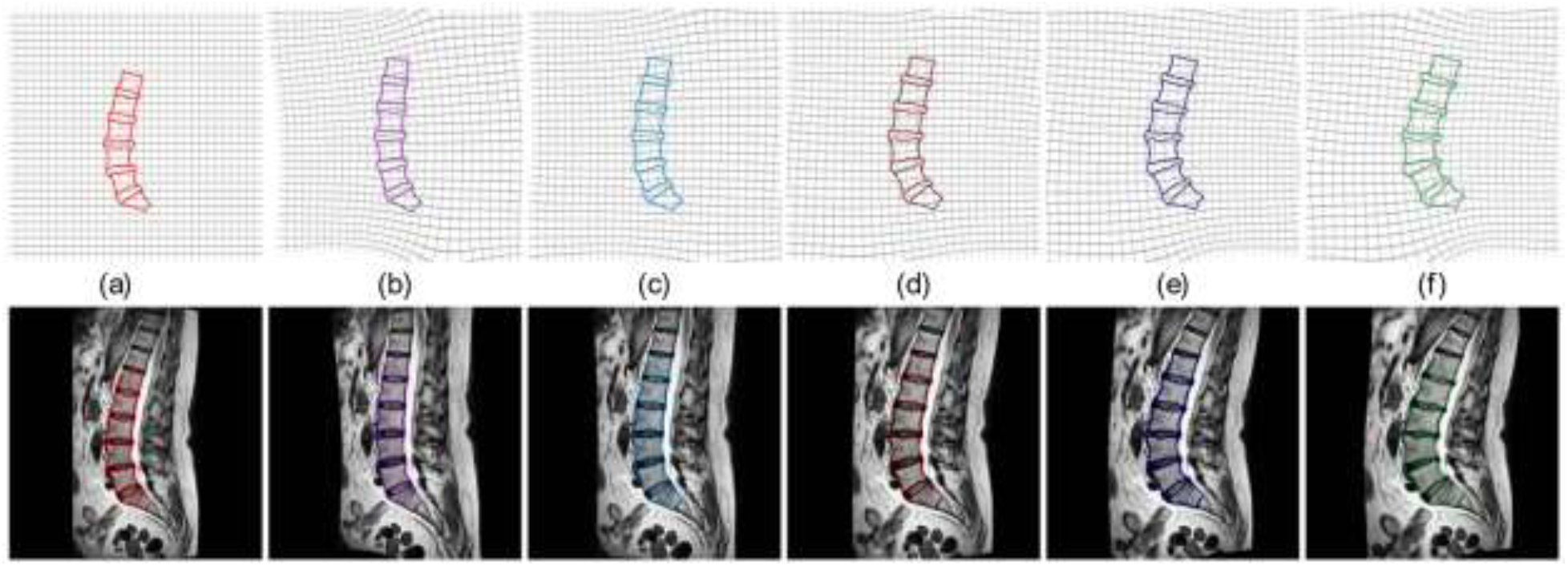
data augmentation/synthesis examples (a-f) using our method. Please zoom in for better visualization.

In the implementation, we obtain the spatial transform *T* from 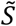 to *S*, and apply the spatial transform *T* to a regular mesh grid around the shape 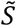 to obtain a deformed grid in the space of the reference image *I*, and then *Ĩ* is obtained by interpolating pixel values of *I* at each node of the deformed grid, i.e., *Ĩ* (*x, y*) = *I*(*T*(*x, y*)) where (*x, y*) denotes a 2D spatial point and *T*(*x, y*) is the transformed point. By using biomechanics and finite element analysis (FEA), the spatial transform, i.e., the deformation field on the mesh grid, is obtained by minimizing the following energy/loss function Π:

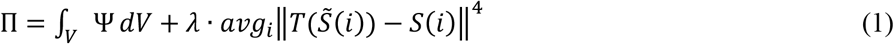

In the above Eq.(1), *V* represents the undeformed mesh grid of the image *Ĩ* to be generated, 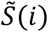 represents the *i*-th point location of the shape 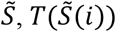 is the transformed point location that needs to be equal to *S*(*i*), and ‖▪‖ denotes vector L2 norm. *avg* is the average operator. *λ* is a weight constant (set to 16 in experiments). Ψ is the strain energy density function that is determined by deformation and mechanical property of soft biological tissues around the lumbar spine. From the perspective of FEA and biomechanics, the Eq.(1) simulates the scenario that under the external “force” proportional to 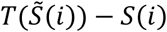 at each point of the lumbar spine shape, the soft biological tissues of human body will deform and reach to an equilibrium state of minimum energy. To speed up the optimization process, we use a deep neural network with sine activation functions to parameterize the transform *T*, i.e., *T*(*x, y*) = *DNN*(*x, y*), and then the energy function in Eq.(1) becomes a function of the DNN internal parameters. The energy optimization problem is resolved by adjusting/optimizing the parameters of the DNN. Once the optimization is done, the deformation field is obtained and then the image *Ĩ* is generated. Since our goal is to generate plausible images for model training and evaluation in the image segmentation tasks, not for patient-specific FEA simulation of human body deformation, we made an assumption about the strain energy density function to reduce computation cost: tissue mechanical behavior follows the Ogden hyperelastic model with homogeneous tissue properties (Ogden and Hill, 1997; Dwivedi et al., 2022). The whole procedure is implemented by using our newly developed PyTorch-FEA library for large deformation biomechanics (Liang et al., 2023).

### 3.2. Loss function

For each model, we use the original loss function if it applicable to our application. If the original loss function is not suitable (e.g., it is for binary classification only), then we use the loss function *ℒ* that combines a Dice loss *ℒ*_*Dice*_ and an area-weighted cross entropy loss *ℒ*_*aw*_*ce*_.

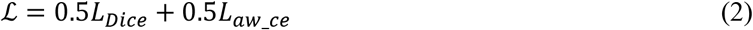

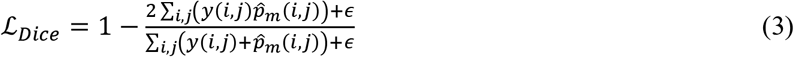

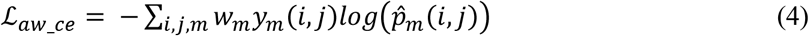

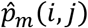 is the m-th element in the output tensor from the softmax layer at the pixel location (*i, j*), which corresponds to the m-th object (a disc or a vertebra) at the location (*i, j*). *y*(*i, j*) is the true label of the pixel at location (*i, j*). *w*_*m*_ is a nonnegative weight inversely proportional to the area of the m-th object, and ∑_*m*_ *w*_*m*_ = 1. *ϵ* is a small constant (1e-4) to prevent the case of 0/0 in Eq.(3). In a lumbar spine image, where the background area is substantially larger than the combined area of the discs and vertebrae, employing area-weighted cross entropy loss effectively reduces the influence of the background in the loss function.

## 4. Experiments

### 4.1. Original Dataset and Augmented Datasets

Our dataset consists of a total of 100 patients’ lumbar spine MR images from the University of Miami medical school, with personal identification information removed. Following the protocol in (Hu et al., 2018), five lumbar discs (D1, D2, D3, D4, D5) and six vertebral bones (L1, L2, L3, L4, L5, S1) in each patient’s mid-sagittal MR image was manually annotated by three trained operators to identify and mark the boundaries and landmarks of the lumbar discs and vertebrae. To ensure accuracy and consistency, the three operators engaged in discussion to reach a consensus on the best annotation (i.e., ground-truth), for each mid-sagittal MR image. In the literature, lumbar disc D1 is also called L1/L2, similarly D2 for L2/L3, D3 for L3/L4, D4 for L4/L5, and D5 for L5/S1. The MR images are of various resolutions, and each of the images are resized to 512 × 512. Each image is also pre-processed independently by normalizing the intensities into range [0,1]. The average pixel spacing is 0.7004mm. The dataset of 100 patients was divided into 70 training samples, 10 validation samples, and 20 test samples, and those samples are referred to as the original training/validation/test samples in this paper.

Subsequently, the augmented dataset, named SSMSpine, is generated by using our method in Section 3.1. The SSMSpine dataset is divided into three sets: an augmented training set with 7000 samples, an augmented validation set with 250 samples, and an augmented test set with 2500 samples. To generate the augmented test set, an SSM was constructed using the 20 original test samples, and then 125 virtual shapes were generated from the SSM. Using each of the 20 original test samples as a reference image and each of the 125 virtual shapes, 20 × 125 (=2500) new MR images were generated using the method in Section 3.1. The augmented training and validation sets were generated in a similar way using another SSM built on the original training and validation samples.

### 4.2. Model evaluation and comparison

In our study, we trained and compared a total of 15 models on our lumbar spine MRI dataset. These models are Attention U-Net (Oktay et al., 2018), HSNet (Zhang et al., 2022), Inception-SwinUnet (Pu et al., 2023), MedT (Valanarasu et al., 2021), MultiResUNet (Ibtehaz and Rahman, 2020), SLT-Net (Feng et al., 2022), Swin-Unet (Cao et al., 2023), UNETR (Hatamizadeh et al., 2021), Swin UNETR (Hatamizadeh et al., 2022), TransUNet (Chen et al., 2021), UCTransNet (Wang et al., 2022), UNet++ (Zhou et al., 2018), UNeXt (Valanarasu and Patel, 2022), UTNet (Gao et al., 2021), and BianqueNet (Zheng et al., 2022). We were unable to test all models mentioned in Section 2 due to either unavailability (e.g., no source code) or incompatibility (e.g., size not matching). By training and evaluating these models, we aimed to compare their performance and determine the most effective approach for spine MRI instance segmentation on our dataset. To ensure compatibility with our dataset, we made minor adjustments to the original codes of some models if necessary.

We conducted two experiments: experiment-A and experiment-B. In experiment-A, each model is trained using the original training set with elastic deformation (Section 3.1). The top 4 models in experiment-A are selected for training using the augmented training set in experiment-B. In both experiments, we applied random translations to the input images within 16 pixels during training, which is a common data augmentation method. In both experiments, the augmented validation set is used for hyper-parameter tuning. The evaluation process contains instance segmentation evaluation.

In the instance segmentation evaluation, we assess model segmentation performance for individual lumbar spine instances/objects in the MR images. The instance segmentation task is formulated as a task of labeling 12 distinct objects (5 lumbar discs, 6 vertebrae, and a background). The input to each model is a single-channel mid-sagittal lumbar spine MR image with size of 512 × 512 pixels. In the segmentation output, each class is represented by a distinct channel as a binary segmentation map. We employ both the Dice Similarity Coefficient and the 95% Hausdorff Distance (HD95) as evaluation metrics. In both experiments, the original test set with 20 samples and the augmented test set with 2500 samples are used separately for model performance assessment on unseen data.

Each model was trained on a Nvidia A6000 GPU with 48GB VRAM. During the training process, a batch size of 6 was used for most of the models, except for training MedT, where a batch size of 2 was utilized due to its large model size. The Adam optimizer with an initial learning rate of 0.0001 was employed for model optimization. A low learning rate is generally preferred to ensure stable convergence during training. Although a low learning rate might slow down the convergence process, it helps avoid convergence failures. Gradient clipping is applied during training to prevent potentially large gradients from causing instability in the learning process. We performed model selection based on the performance on the validation set.

### 4.3. Results of Experiment-A with the original training set

In experiment-A, the 15 models were trained on the original training set with elastic deformation and random-shift, and then the models were evaluated on both the original test set and the augmented test set to measure instance segmentation accuracy. Figure 3 displays the performance of the top 4 models.

**Fig. 3.**
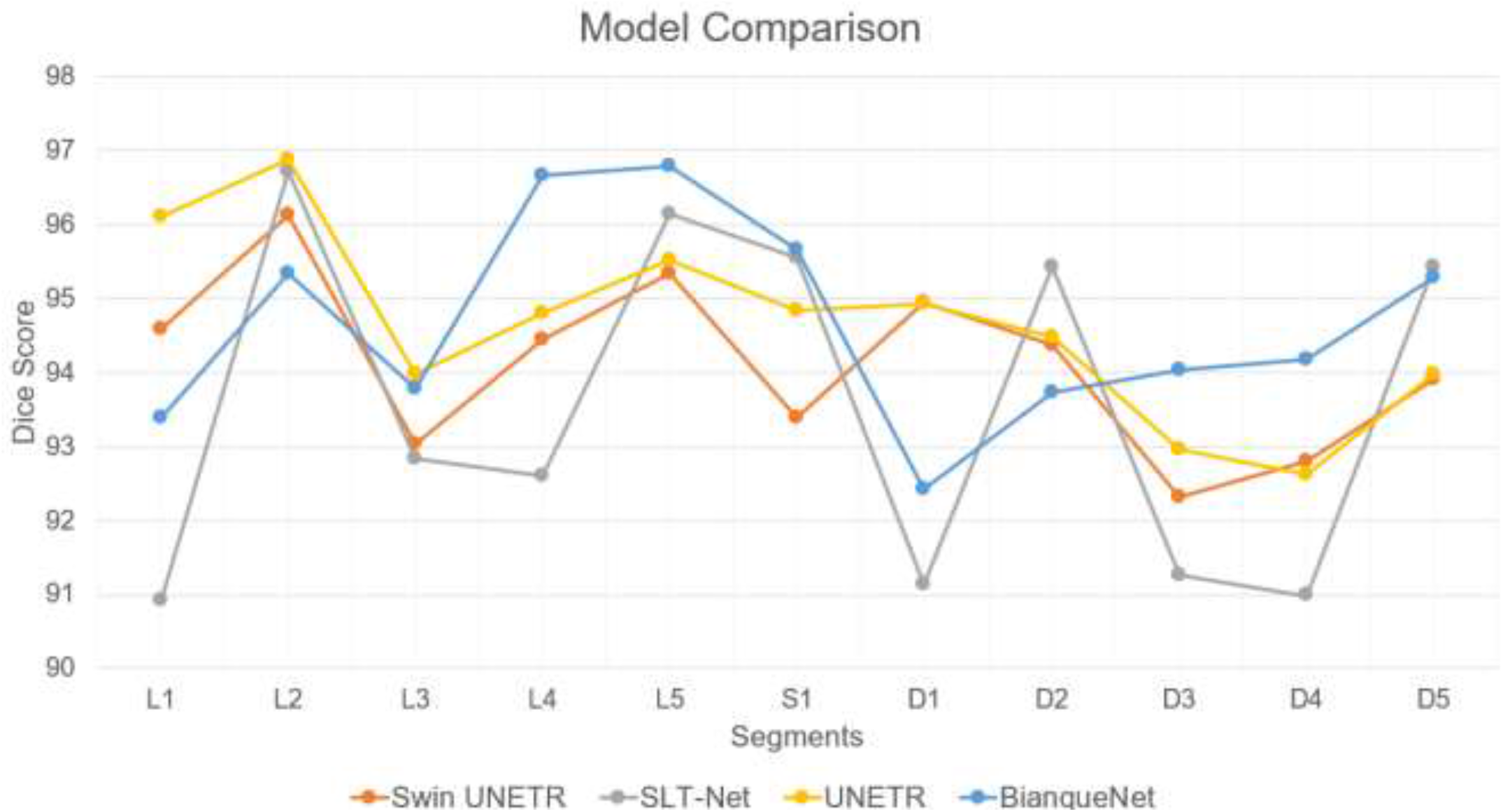
Top 4 Model Comparison Results (Dice) on the augmented test set

#### 4.3.1. Instance Segmentation Evaluation

Table 2 summarizes the instance segmentation results for vertebrae bodies (VB) and intervertebral discs (IVD) in terms of the Dice Similarity Coefficient on the original test set consisting of 20 samples. For better clarity and ease of understanding, we have converted Dice Similarity Coefficient into percentage ratios between 0 and 100%. The results show that Transformer-based models surpasses the CNN-only models.

**Table 2.** Dice (the higher, the better) of each model on the original test set.

|  | L1 | L2 | L3 | L4 | L5 | S1 | D1 | D2 | D3 | D4 | D5 | Average |
| --- | --- | --- | --- | --- | --- | --- | --- | --- | --- | --- | --- | --- |
| Swin UNETR | 93.126<br>±7.356 | 96.813<br>±1.641 | 93.874<br>±11.389 | **95.805**<br>±3.189 | 94.380<br>±4.816 | 92.194<br>±6.449 | 92.757<br>±4.568 | **94.299**<br>±3.519 | 92.821<br>±6.563 | 89.777<br>±7.645 | 91.077<br>±10.179 | **93.357**<br>±4.098 |
| SLT-Net | 91.967<br>±6.991 | 97.052<br>±0.902 | **94.328**<br>±10.261 | 93.619<br>±13.504 | 95.763<br>±2.553 | 93.805<br>±1.815 | 92.990<br>±4.893 | 92.938<br>±6.914 | 90.282<br>±16.896 | 91.481<br>±5.639 | 92.589<br>±6.048 | 93.347<br>±5.886 |
| UNETR | **94.355**<br>±3.131 | 96.795<br>±1.709 | 91.900<br>±18.125 | 95.546<br>±3.196 | 94.099<br>±3.814 | 92.988<br>±4.539 | 93.261<br>±3.878 | 93.554<br>±4.254 | 90.931<br>±9.003 | 90.435<br>±5.678 | 91.446<br>±4.198 | 93.210<br>±4.333 |
| BianqueNet | 89.330<br>±21.689 | 93.332<br>±16.860 | 91.819<br>±18.667 | 95.245<br>±6.709 | **96.011**<br>±2.155 | **94.193**<br>±5.037 | 90.069<br>±21.017 | 92.566<br>±9.930 | **93.635**<br>±4.852 | **93.539**<br>±3.373 | **94.326**<br>±4.325 | 93.097<br>±2.840 |
| Inception-SwinUnet | 91.397<br>±19.415 | **97.335**<br>±0.812 | 91.926<br>±20.366 | 93.837<br>±12.637 | 91.156<br>±18.678 | 91.592<br>±8.410 | **93.533**<br>±4.614 | 93.281<br>±7.055 | 89.739<br>±20.753 | 87.727<br>±14.249 | 91.233<br>±11.216 | 92.069<br>±9.514 |
| HSNet | 94.240<br>±12.384 | 94.116<br>±10.544 | 90.123<br>±22.355 | 91.649<br>±21.243 | 93.456<br>±11.688 | 92.662<br>±9.901 | 92.948<br>±7.903 | 88.941<br>±21.43 | 89.008<br>±21.266 | 89.158<br>±17.099 | 91.986<br>±12.085 | 91.662<br>±13.085 |
| Swin-Unet | 90.839<br>±20.912 | 93.514<br>±16.261 | 91.781<br>±21.085 | 91.781<br>±21.08 | 90.076<br>±21.037 | 92.197<br>±6.612 | 91.638<br>±11.014 | 90.000<br>±20.632 | 89.755<br>±20.720 | 87.864<br>±20.629 | 88.554<br>±16.934 | 90.727<br>±16.984 |
| UNeXt | 92.354<br>±15.512 | 95.835<br>±4.808 | 91.220<br>±21.036 | 89.282<br>±21.93 | 88.818<br>±18.828 | 87.325<br>±14.428 | 92.082<br>±9.072 | 91.271<br>±16.774 | 87.280<br>±21.834 | 85.790<br>±19.574 | 89.065<br>±17.579 | 90.029<br>±12.849 |
| TransUNet | 92.245<br>±9.199 | 87.526<br>±25.59 | 88.178<br>±24.371 | 91.605<br>±21.122 | 91.219<br>±21.025 | 89.705<br>±20.496 | 81.444<br>±32.928 | 86.309<br>±24.176 | 88.812<br>±21.076 | 88.060<br>±20.596 | 90.294<br>±20.782 | 88.672<br>±20.293 |
| MedT | 88.707<br>±20.984 | 94.461<br>±8.57 | 91.011<br>±20.989 | 90.999<br>±20.981 | 87.121<br>±25.111 | 87.183<br>±15.317 | 87.639<br>±20.944 | 89.689<br>±19.384 | 87.318<br>±20.279 | 82.663<br>±23.855 | 85.180<br>±22.131 | 88.361<br>±18.641 |
| UTNet | 81.322<br>±33.18 | 82.886<br>±31.442 | 85.159<br>±29.06 | 88.802<br>±24.539 | 90.776<br>±20.988 | 87.080<br>±14.823 | 77.511<br>±32.691 | 78.711<br>±32.650 | 83.856<br>±28.478 | 85.608<br>±21.674 | 89.312<br>±18.307 | 84.639<br>±23.716 |
| UCTransNet | 75.346<br>±31.945 | 78.997<br>±28.28 | 81.862<br>±26.04 | 80.907<br>±29.371 | 86.107<br>±22.629 | 86.968<br>±20.202 | 80.058<br>±25.635 | 80.407<br>±22.833 | 76.787<br>±30.266 | 81.144<br>±25.322 | 88.063<br>±19.596 | 81.513<br>±20.303 |
| Attention U-Net | 74.333<br>±35.642 | 74.288<br>±33.164 | 68.732<br>±36.905 | 76.499<br>±29.188 | 86.964<br>±22.583 | 85.474<br>±25.111 | 73.700<br>±37.113 | 68.943<br>±37.476 | 68.092<br>±35.719 | 80.147<br>±25.685 | 87.694<br>±21.912 | 76.806<br>±25.335 |
| UNet++ | 68.734<br>±40.68 | 66.998<br>±35.694 | 74.033<br>±27.911 | 71.468<br>±40.229 | 81.297<br>±31.709 | 83.980<br>±27.527 | 65.450<br>±39.937 | 71.473<br>±28.932 | 67.208<br>±40.117 | 79.834<br>±28.708 | 84.529<br>±26.387 | 74.091<br>±26.461 |
| MultiResUNet | 77.421<br>±31.596 | 74.309<br>±29.915 | 66.317<br>±34.739 | 66.828<br>±35.945 | 76.870<br>±35.698 | 84.644<br>±22.624 | 75.710<br>±33.508 | 70.025<br>±27.873 | 58.563<br>±39.241 | 72.553<br>±33.342 | 75.217<br>±31.98 | 72.587<br>±26.547 |

Table 3 shows the instance segmentation results measured by 95% Hausdorff distance (HD95) on the original test set. The results show that Transformer-based models surpasses the CNN-only models.

**Table 3.** HD95 (the lower, the better) of each model on the original test set.

|  | L1 | L2 | L3 | L4 | L5 | S1 | D1 | D2 | D3 | D4 | D5 | Average |
| --- | --- | --- | --- | --- | --- | --- | --- | --- | --- | --- | --- | --- |
| Swin UNETR | 8.594<br>±12.047 | 11.024<br>±33.718 | 2.992<br>±3.483 | 3.147<br>±6.521 | 3.759<br>±5.905 | 9.454<br>±13.059 | 19.057<br>±41.678 | 1.894<br>±2.472 | 3.466<br>±5.801 | 3.814<br>±3.828 | 8.128<br>±12.917 | 6.848<br>±10.137 |
| SLT-Net | 13.716<br>±16.122 | 1.332<br>±0.64 | 2.371<br>±3.562 | 2.778<br>±5.493 | 2.363<br>±2.252 | 8.11<br>±17.93 | 2.348<br>±3.912 | 2.004<br>±2.338 | 3.118<br>±5.032 | 2.865<br>±2.174 | 4.762<br>±12.584 | 4.161<br>±4.089 |
| UNETR | 3.744<br>±6.201 | 1.512<br>±1.261 | 3.679<br>±7.65 | 2.237<br>±1.722 | 5.927<br>±6.302 | **4.724**<br>±5.909 | 2.442<br>±3.25 | **1.704**<br>±1.3 | 3.589<br>±4.982 | 3.277<br>±2.968 | 7.517<br>±8.093 | 3.668<br>±2.34 |
| BianqueNet | 5.056<br>±10.105 | 7.158<br>±15.206 | 3.624<br>±7.993 | 3.375<br>±7.988 | **2.290**<br>±2.285 | 5.308<br>±9.319 | 4.278<br>±9.886 | 5.106<br>±10.580 | 3.138<br>±7.382 | 5.857<br>±12.764 | **3.713**<br>±6.602 | 4.446<br>±9.739 |
| Inception-SwinUnet | 3.378<br>±7.275 | **1.197**<br>±0.555 | 3.5<br>±8.067 | 3.033<br>±6.334 | 3.362<br>±3.809 | 6.39<br>±10.21 | 7.53<br>±25.967 | 2.051<br>±3.129 | **1.59**<br>±0.876 | 3.957<br>±4.714 | 5.715<br>±9.957 | 3.791<br>±4.134 |
| HSNet | 3.006<br>±6.714 | 3.634<br>±8.238 | 4.121<br>±8.874 | 3.94<br>±9.34 | 5.547<br>±11.36 | 5.196<br>±10.482 | 2.869<br>±7.442 | 5.378<br>±11.527 | 4.145<br>±9.551 | 5.966<br>±11.862 | 5.045<br>±11.035 | 4.441<br>±7.376 |
| Swin-Unet | **2.131**<br>±3.459 | 2.226<br>±4.686 | 1.753<br>±1.591 | **1.971**<br>±2.484 | 2.767<br>±2.596 | 4.732<br>±7.47 | **2.221**<br>±3.907 | 2.937<br>±6.313 | 1.679<br>±1.159 | **2.08**<br>±1.308 | 6.339<br>±10.19 | **2.803**<br>±2.24 |
| UNeXt | 2.805<br>±4.959 | 2.016<br>±2.768 | **1.933**<br>±1.542 | 7.259<br>±19.018 | 13.485<br>±17.479 | 10.8<br>±15.286 | 5.98<br>±14.131 | 2.536<br>±4.028 | 5.69<br>±15.364 | 7.292<br>±13.081 | 11.01<br>±15.926 | 6.437<br>±8.162 |
| TransUNet | 7.727<br>±15.874 | 8.556<br>±20.98 | 9.261<br>±19.824 | 3.836<br>±9.163 | 4.372<br>±9.063 | 7.595<br>±12.805 | 8.968<br>±22.198 | 8.662<br>±19.145 | 3.876<br>±9.056 | 5.176<br>±10.127 | 4.15<br>±9.494 | 6.562<br>±11.646 |
| MedT | 8.665<br>±14.016 | 2.557<br>±2.564 | 6.168<br>±11.39 | 2.386<br>±2.751 | 7.056<br>±10.149 | 10.406<br>±10.254 | 5.684<br>±10.784 | 7.024<br>±11.272 | 63.762<br>±83.727 | 5.359<br>±8.133 | 18.524<br>±33.337 | 12.508<br>±7.994 |
| UTNet | 6.897<br>±12.365 | 7.209<br>±12.726 | 10.986<br>±16.485 | 5.092<br>±10.427 | 8.523<br>±13.468 | 20.203<br>±35.303 | 7.421<br>±13.484 | 13.133<br>±18.251 | 11.486<br>±17.583 | 10.463<br>±17.173 | 8.903<br>±13.608 | 10.029<br>±12.359 |
| UCTransNet | 11.554<br>±13.053 | 21.176<br>±20.644 | 30.833<br>±45.698 | 12.752<br>±15.773 | 11.034<br>±15.015 | 9.536<br>±15.319 | 20.424<br>±32.647 | 23.576<br>±27.95 | 20.578<br>±22.595 | 14.814<br>±16.837 | 7.444<br>±13.146 | 16.702<br>±12.286 |
| Attention U-Net | 9.394<br>±12.866 | 13.821<br>±17.43 | 16.946<br>±18.222 | 21.225<br>±32.217 | 10.944<br>±15.876 | 7.474<br>±13.279 | 13.878<br>±18.345 | 15.805<br>±19.732 | 28.632<br>±35.708 | 10.027<br>±15.161 | 7.124<br>±13.263 | 14.116<br>±13.022 |
| UNet++ | 6.733<br>±11.95 | 16.872<br>±17.754 | 22.11<br>±20.232 | 15.901<br>±19.52 | 12.515<br>±16.298 | 9.518<br>±14.037 | 9.862<br>±14.455 | 21.599<br>±20.421 | 19.863<br>±26.463 | 10.369<br>±16.036 | 9.929<br>±15.288 | 14.116<br>±11.826 |
| MultiResUNet | 8.841<br>±11.614 | 17.292<br>±16.037 | 19.035<br>±17.013 | 15.814<br>±15.802 | 9.93<br>±13.24 | 9.582<br>±13.03 | 9.63<br>±14.817 | 18.331<br>±18.619 | 22.223<br>±19.32 | 10.918<br>±15.737 | 10.632<br>±14.034 | 13.839<br>±11.524 |

We also assessed the segmentation performance of all models on the augmented test set consisting of 2500 samples. Table 4 (Dice) and Table 5 (HD95) summarize the instance segmentation performance of each model evaluated on the augmented test set.

**Table 4.**
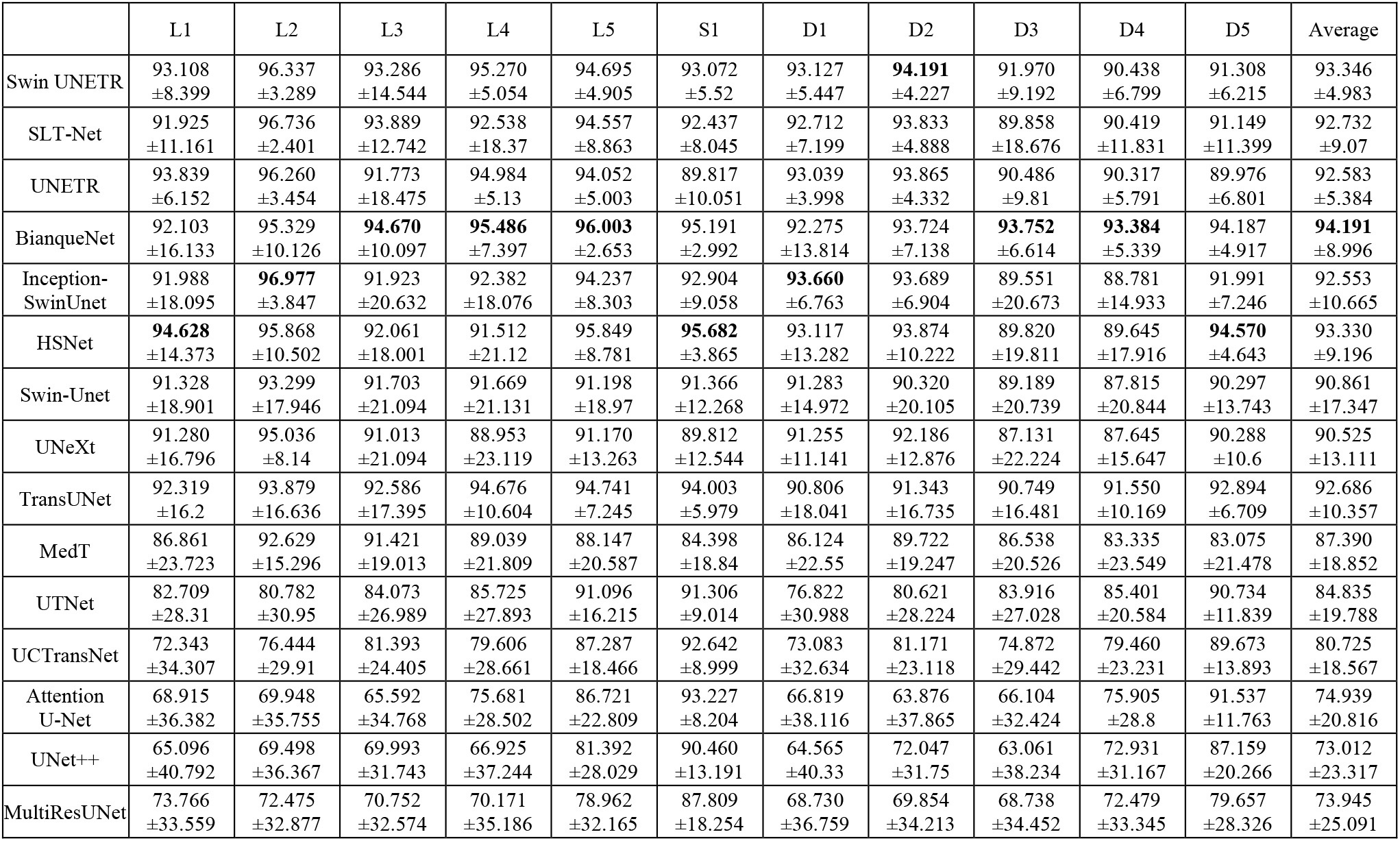
Dice (the higher, the better) of each model on the augmented test set.

|  | L1 | L2 | L3 | L4 | L5 | S1 | D1 | D2 | D3 | D4 | D5 | Average |
| --- | --- | --- | --- | --- | --- | --- | --- | --- | --- | --- | --- | --- |
| Swin UNETR | 93.108<br>±8.399 | 96.337<br>±3.289 | 93.286<br>±14.544 | 95.270<br>±5.054 | 94.695<br>±4.905 | 93.072<br>±5.52 | 93.127<br>±5.447 | **94.191**<br>±4.227 | 91.970<br>±9.192 | 90.438<br>±6.799 | 91.308<br>±6.215 | 93.346<br>±4.983 |
| SLT-Net | 91.925<br>±11.161 | 96.736<br>±2.401 | 93.889<br>±12.742 | 92.538<br>±18.37 | 94.557<br>±8.863 | 92.437<br>±8.045 | 92.712<br>±7.199 | 93.833<br>±4.888 | 89.858<br>±18.676 | 90.419<br>±11.831 | 91.149<br>±11.399 | 92.732<br>±9.07 |
| UNETR | 93.839<br>±6.152 | 96.260<br>±3.454 | 91.773<br>±18.475 | 94.984<br>±5.13 | 94.052<br>±5.003 | 89.817<br>±10.051 | 93.039<br>±3.998 | 93.865<br>±4.332 | 90.486<br>±9.81 | 90.317<br>±5.791 | 89.976<br>±6.801 | 92.583<br>±5.384 |
| BianqueNet | 92.103<br>±16.133 | 95.329<br>±10.126 | **94.670**<br>±10.097 | **95.486**<br>±7.397 | **96.003**<br>±2.653 | 95.191<br>±2.992 | 92.275<br>±13.814 | 93.724<br>±7.138 | **93.752**<br>±6.614 | **93.384**<br>±5.339 | 94.187<br>±4.917 | **94.191**<br>±8.996 |
| Inception-SwinUnet | 91.988<br>±18.095 | **96.977**<br>±3.847 | 91.923<br>±20.632 | 92.382<br>±18.076 | 94.237<br>±8.303 | 92.904<br>±9.058 | **93.660**<br>±6.763 | 93.689<br>±6.904 | 89.551<br>±20.673 | 88.781<br>±14.933 | 91.991<br>±7.246 | 92.553<br>±10.665 |
| HSNet | **94.628**<br>±14.373 | 95.868<br>±10.502 | 92.061<br>±18.001 | 91.512<br>±21.12 | 95.849<br>±8.781 | **95.682**<br>±3.865 | 93.117<br>±13.282 | 93.874<br>±10.222 | 89.820<br>±19.811 | 89.645<br>±17.916 | **94.570**<br>±4.643 | 93.330<br>±9.196 |
| Swin-Unet | 91.328<br>±18.901 | 93.299<br>±17.946 | 91.703<br>±21.094 | 91.669<br>±21.131 | 91.198<br>±18.97 | 91.366<br>±12.268 | 91.283<br>±14.972 | 90.320<br>±20.105 | 89.189<br>±20.739 | 87.815<br>±20.844 | 90.297<br>±13.743 | 90.861<br>±17.347 |
| UNeXt | 91.280<br>±16.796 | 95.036<br>±8.14 | 91.013<br>±21.094 | 88.953<br>±23.119 | 91.170<br>±13.263 | 89.812<br>±12.544 | 91.255<br>±11.141 | 92.186<br>±12.876 | 87.131<br>±22.224 | 87.645<br>±15.647 | 90.288<br>±10.6 | 90.525<br>±13.111 |
| TransUNet | 92.319<br>±16.2 | 93.879<br>±16.636 | 92.586<br>±17.395 | 94.676<br>±10.604 | 94.741<br>±7.245 | 94.003<br>±5.979 | 90.806<br>±18.041 | 91.343<br>±16.735 | 90.749<br>±16.481 | 91.550<br>±10.169 | 92.894<br>±6.709 | 92.686<br>±10.357 |
| MedT | 86.861<br>±23.723 | 92.629<br>±15.296 | 91.421<br>±19.013 | 89.039<br>±21.809 | 88.147<br>±20.587 | 84.398<br>±18.84 | 86.124<br>±22.55 | 89.722<br>±19.247 | 86.538<br>±20.526 | 83.335<br>±23.549 | 83.075<br>±21.478 | 87.390<br>±18.852 |
| UTNet | 82.709<br>±28.31 | 80.782<br>±30.95 | 84.073<br>±26.989 | 85.725<br>±27.893 | 91.096<br>±16.215 | 91.306<br>±9.014 | 76.822<br>±30.988 | 80.621<br>±28.224 | 83.916<br>±27.028 | 85.401<br>±20.584 | 90.734<br>±11.839 | 84.835<br>±19.788 |
| UCTransNet | 72.343<br>±34.307 | 76.444<br>±29.91 | 81.393<br>±24.405 | 79.606<br>±28.661 | 87.287<br>±18.466 | 92.642<br>±8.999 | 73.083<br>±32.634 | 81.171<br>±23.118 | 74.872<br>±29.442 | 79.460<br>±23.231 | 89.673<br>±13.893 | 80.725<br>±18.567 |
| Attention U-Net | 68.915<br>±36.382 | 69.948<br>±35.755 | 65.592<br>±34.768 | 75.681<br>±28.502 | 86.721<br>±22.809 | 93.227<br>±8.204 | 66.819<br>±38.116 | 63.876<br>±37.865 | 66.104<br>±32.424 | 75.905<br>±28.8 | 91.537<br>±11.763 | 74.939<br>±20.816 |
| UNet++ | 65.096<br>±40.792 | 69.498<br>±36.367 | 69.993<br>±31.743 | 66.925<br>±37.244 | 81.392<br>±28.029 | 90.460<br>±13.191 | 64.565<br>±40.33 | 72.047<br>±31.75 | 63.061<br>±38.234 | 72.931<br>±31.167 | 87.159<br>±20.266 | 73.012<br>±23.317 |
| MultiResUNet | 73.766<br>±33.559 | 72.475<br>±32.877 | 70.752<br>±32.574 | 70.171<br>±35.186 | 78.962<br>±32.165 | 87.809<br>±18.254 | 68.730<br>±36.759 | 69.854<br>±34.213 | 68.738<br>±34.452 | 72.479<br>±33.345 | 79.657<br>±28.326 | 73.945<br>±25.091 |

**Table 5.**
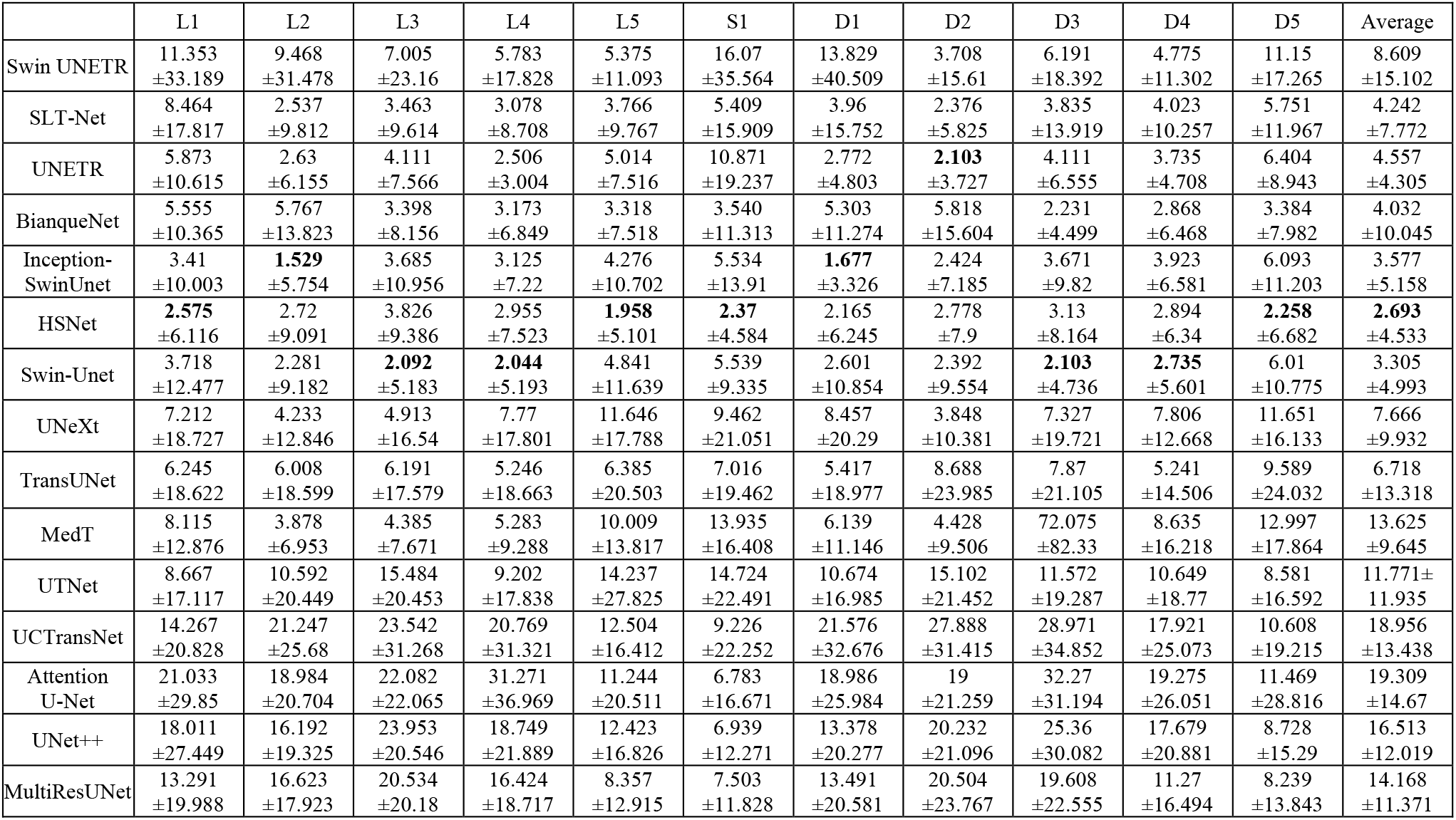
HD95 value (the lower, the better) of each model on the augmented test set.

Figure 4 shows segmentation examples of the 15 models. All of the models produce misclassifications or fragmentation errors. For example, BianqueNet exhibits hollow holes in both D3 and D4. TransUnet incorrectly identifies D3 as D2 and exhibits a significant segmentation error in the lower left corner in the MR scan. Also seen in Figure 4, many models produce incorrect predictions around pixels in close proximity to the boundary of two adjacent objects.

**Fig. 4.**
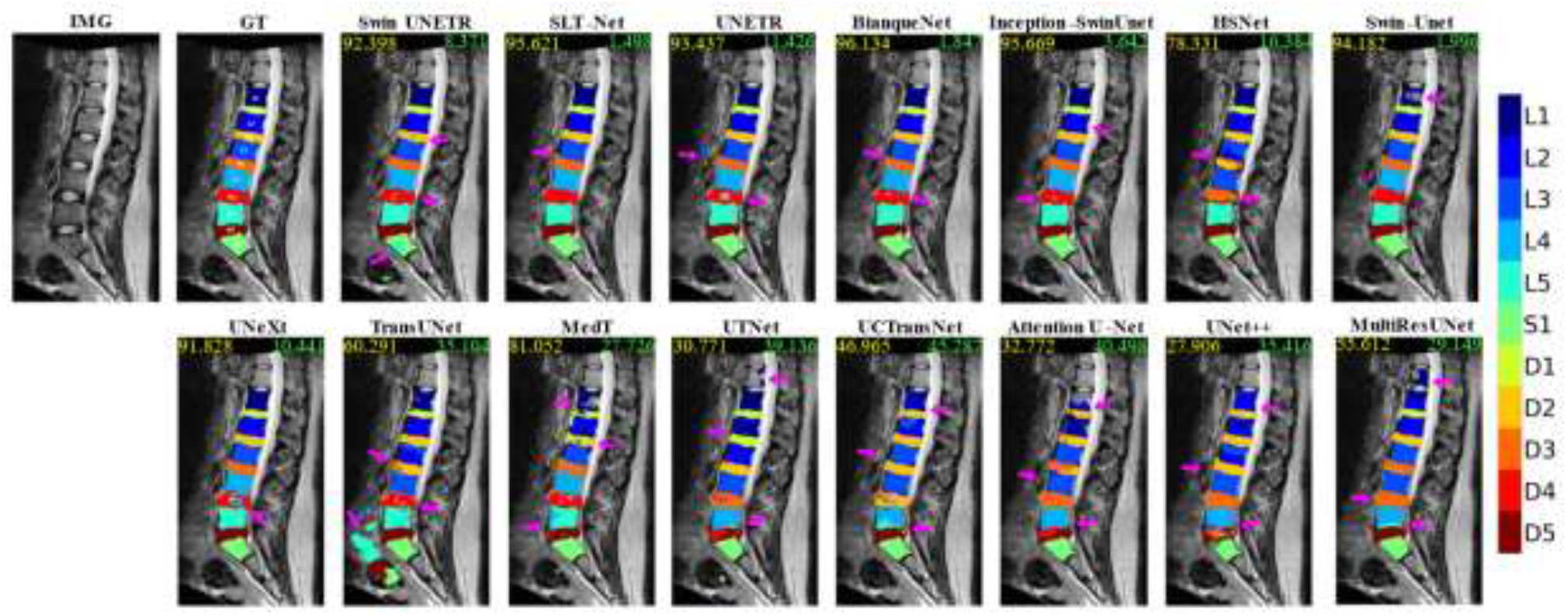
Segmentation examples of the 15 models. IMG is the input image. GT indicates the Ground-Truth annotation. Dice (green color) and HD95 (yellow color) are shown on top of each image. The 11 lumbar objects are shown in different colors. Segmenttion errors are indicated by pink arrows.

### 4.4. Results of Experiment-B with the augmented training set

In this section, we show the advantages of our data augmentation method in Section 3.1. The top four models in Table 2 are Swin UNETR, SLT-Net, UNETR, BianqueNet. In the experiment-B, each of the four models was trained from scratch using the augmented training set consisting of 7000 samples. Model evaluations were conducted on the augmented test set (see Table 8 and Table 9). The results show that the segmentation performance of most models is improved, suggesting that the augmented data can enhance the generalization and overall performance of a model.

**Table 8.** Dice (the higher, the better) of each model on the augmented test set. The “change” is the Dice difference between training on the augmented training set and training on the original training set.

|  | L1 | L2 | L3 | L4 | L5 | S1 | D1 | D2 | D3 | D4 | D5 | Average | Change |
| --- | --- | --- | --- | --- | --- | --- | --- | --- | --- | --- | --- | --- | --- |
| Swin UNETR | 94.593<br>±8.113 | 96.124<br>±7.352 | 93.027<br>±17.824 | 94.442<br>±12.723 | 95.345<br>±6.840 | 93.384<br>±6.406 | 94.949<br>±3.461 | 94.369<br>±7.674 | 92.323<br>±13.660 | 92.806<br>±5.711 | 93.900<br>±4.372 | 94.115<br>±9.581 | **+0.769** |
| SLT-Net | 90.928<br>±21.398 | 96.708<br>±5.097 | 92.836<br>±18.462 | 92.604<br>±19.937 | 96.143<br>±4.279 | 95.545<br>±2.669 | 91.135<br>±18.062 | 95.435<br>±3.448 | 91.254<br>±17.130 | 90.982<br>±15.676 | 95.427<br>±2.972 | 93.545<br>±14.102 | **+0.813** |
| UNETR | 96.099<br>±2.574 | 96.878<br>±2.479 | 93.981<br>±11.711 | 94.798<br>±8.922 | 95.509<br>±3.888 | 94.832<br>±2.951 | 94.930<br>±2.159 | 94.486<br>±5.341 | 92.957<br>±8.036 | 92.616<br>±4.935 | 93.974<br>±3.926 | 94.642<br>±6.081 | **+2.059** |
| BianqueNet | 93.381<br>±16.166 | 95.339<br>±12.916 | 93.786<br>±13.760 | 96.670<br>±4.308 | 96.789<br>±2.243 | 95.666<br>±2.302 | 92.427<br>±16.211 | 93.723<br>±11.633 | 94.040<br>±7.633 | 94.184<br>±4.318 | 95.276<br>±3.268 | 94.662<br>±10.231 | **+0.471** |

**Table 9.** HD95 (the lower, the better) of each model on the augmented test set. The “change” is the HD95 difference between training on the augmented training set and training on the original training set.

|  | L1 | L2 | L3 | L4 | L5 | S1 | D1 | D2 | D3 | D4 | D5 | Average | Change |
| --- | --- | --- | --- | --- | --- | --- | --- | --- | --- | --- | --- | --- | --- |
| Swin UNETR | 17.591<br>±52.428 | 13.865<br>±44.911 | 12.207<br>±42.917 | 11.290<br>±38.870 | 10.393<br>±35.903 | 19.144<br>±52.802 | 9.570<br>±35.018 | 12.249<br>±43.617 | 7.145<br>±30.825 | 5.524<br>±21.039 | 11.808<br>±30.508 | 11.889<br>±40.236 | **+3.280** |
| SLT-Net | 3.720<br>±8.774 | 1.577<br>±3.310 | 3.227<br>±7.838 | 1.899<br>±5.205 | 2.012<br>±2.264 | 1.871<br>±2.843 | 2.344<br>±6.312 | **1.387**<br>±2.514 | 3.100<br>±8.190 | 2.584<br>±4.752 | 1.602<br>±2.714 | 2.302<br>±5.549 | **-1.940** |
| UNETR | 1.923<br>±2.598 | 1.774<br>±3.164 | 2.679<br>±4.416 | 3.029<br>±7.859 | 3.056<br>±5.502 | 3.340<br>±5.462 | 1.329<br>±1.5817 | 1.788<br>±2.9369 | 2.348<br>±3.860 | 2.640<br>±3.108 | 2.455<br>±3.079 | 2.396<br>±4.341 | **-2.161** |
| BianqueNet | 3.815<br>±9.985 | 2.907<br>±8.092 | 3.849<br>±10.217 | 2.605<br>±9.389 | 2.268<br>±7.335 | 2.667<br>±10.589 | 3.383<br>±9.089 | 4.257<br>±11.508 | 3.149<br>±10.065 | 2.061<br>±5.126 | 2.860<br>±13.087 | 3.075<br>±9.734 | **-0.957** |

It is noteworthy that training Transformer-based models with limited data is challenging. Nevertheless, the new data augmentation method effectively tackles the challenge of data scarcity with the support of SSM and biomechanics. Our data augmentation can generate synthetic images that closely resemble real data, effectively enhancing the model training process. This approach assists in augmenting the available data, enabling Transformer models to learn from a more diverse and representative dataset. The augmented/synthesized datasets can be made publicly available without any concerns related to medical data privacy.

By using the instance segmentation evaluation, a comprehensive evaluation of each model’s performance is achieved. This ensures that models excel not only instance segmentation ability but also maintain their accuracy under realistic and varied conditions.

#### 4.4.1. Instance Segmentation Evaluation

Table 8 (Dice) and Table 9 (HD95) present the instance segmentation evaluation results on the augmented test set. It is evident that the segmentation performances of all models have improved by using the augmented dataset, as indicated by enhancements in both evaluation metrics, except for Swin UNETR on the HD95 metric. The augmented dataset offers a broader domain, enabling the models to acquire more diverse knowledge and achieve improved segmentation performance. This increased resilience to data variations is a notable benefit of the augmented training dataset. Increasing the complexity of the training dataset helps prevent overfitting and regulating models from memorizing the training.

We further analyzed the distribution of the dice scores of each model by calculating minimum (min) and the 5^th^ percentile (Q5) of the dice scores across the samples in the augmented test set. The results are reported in Table 10, which shows that all of the models make huge mistakes on some samples in the test set.

**Table 10.** min and Q5 of Dice scores of each model on the augmented test set.

|  | Swin_UNETR |  | SLT_Net |  | UNETR |  | BianqueNet |  |
| --- | --- | --- | --- | --- | --- | --- | --- | --- |
|  | min | Q5 | min | Q5 | min | Q5 | min | Q5 |
| L1 | 14.439 | 81.683 | 0.000 | 26.034 | 76.269 | 90.137 | 0.000 | 81.142 |
| L2 | 9.578 | 88.879 | 52.846 | 94.887 | 76.976 | 92.733 | 0.000 | 93.437 |
| L3 | 0.000 | 85.303 | 0.000 | 83.608 | 35.331 | 86.860 | 0.000 | 66.823 |
| L4 | 0.000 | 87.625 | 0.000 | 64.281 | 45.150 | 83.110 | 0.549 | 91.171 |
| L5 | 11.825 | 88.599 | 0.000 | 91.660 | 58.809 | 87.756 | 33.610 | 93.513 |
| S1 | 40.717 | 78.308 | 0.000 | 92.101 | 38.942 | 89.988 | 69.128 | 91.717 |
| D1 | 64.113 | 87.755 | 0.000 | 73.123 | 85.688 | 89.586 | 0.000 | 81.372 |
| D2 | 18.270 | 84.046 | 60.219 | 91.264 | 57.125 | 90.432 | 0.000 | 85.795 |
| D3 | 0.000 | 72.141 | 0.000 | 74.561 | 51.413 | 75.504 | 1.579 | 83.127 |
| D4 | 29.136 | 82.262 | 0.000 | 83.406 | 63.937 | 82.084 | 0.000 | 89.208 |
| D5 | 61.554 | 83.589 | 4.772 | 91.999 | 69.841 | 83.746 | 55.854 | 88.920 |
| whole | 25.071 | 86.539 | 36.613 | 74.175 | 67.425 | 87.993 | 14.611 | 86.377 |

## 5. Conclusion

In this paper, we present an evaluation of 15 DNN models for instance segmentation of lumbar spine MR images, using the original dataset of 100 patients and the generated SSMSpine dataset of thousands of virtual patients. We developed the SSM-biomechanics based data augmentation method to further improve model performance by providing large and diverse datasets of synthetic images with ground-truth. Given that our augmented datasets consist entirely of synthetic data, we have made our augmented dataset, SSMSpine, publicly available. The results presented indicate that models trained on the augmented training set had comparably or even better performance than the same models trained on the original training set. This underscores that our data augmentation method can generate synthetic data that eliminates privacy concerns while retaining in the same image domain.

Our current study mainly focused on the mid-sagittal lumbar spine MR images for two major reasons. First, as shown in a clinical study (Hu et al., 2018), the mid-sagittal image of a patient provides the most useful information for the diagnosis of lumbar spine degeneration. Second, the slice thickness of a lumbar MR scan in the sagittal direction is often much larger than 5mm, which causes difficulties to create accurate 3D ground-truth annotation for model training. Nevertheless, the models could be directly extended to handle 3D images once the slice thickness becomes acceptably small with the advancement of imaging technology.

The 15 DNN models, including the Transformer-based models, often generate segmentation artifacts, such as: (1) extra areas not belonging to any discs or vertebrae, (2) assigning the same class label to two different discs, and (3) broken area of a disc or vertebrae. Thus, new models are needed for artifact-free geometry reconstruction of lumbar spine from MR images.

